# Early induction of antibacterial activities distinguishes response of mice to infection with non-permissive from response to permissive *Salmonella*

**DOI:** 10.1101/2020.12.07.414268

**Authors:** Jitender Yadav, Ayub Qadri

## Abstract

*Salmonella enterica* serovar Typhi (*S*. Typhi), the causative agent of typhoid in humans, shares very high homology with closely related serovar, *S*. Typhimurium. Yet, unlike *S*. Typhimurium, *S*. Typhi does not establish infection in mice, the reasons for which are not well understood. Here, we present evidence that unlike *S*. Typhimurium, *S*. Typhi brings about induction of extracellular and intracellular antibacterial activities in mice. Cell-free peritoneal fluids from *S*. Typhi but not *S*. Typhimurium-infected mice inhibited replication of *Salmonella ex vivo*. The production of this activity was reduced in presence of serine protease inhibitor, phenylmethylsulfonlyl fluoride (PMSF). PMSF also inhibited generation of antibacterial activity released from *in vitro S*. Typhi - infected peritoneal macrophages in a cell death – dependent manner. Intracellularly, infection with *S*. Typhi *in vitro* as well as *in vivo* resulted in increased mRNA levels of iron-regulating molecules, ferroportin and lipocalin. These results suggest that induction of antibacterial molecules early on in mice in response to *S*. Typhi may prevent establishment of infection with this *Salmonella* serovar.

## INTRODUCTION

*S*. Typhi produces systemic infection, typhoid, almost exclusively in humans. Only in chimpanzees, infection with a very high inoculum (1 × 10^11^ CFU) of this pathogen produces a disease that resembles human typhoid [1]. Due to this extreme host specificity, there is no animal model for *S*. Typhi infection because of which pathogenesis of human typhoid, host-pathogen interactions that ensue during infection with *S*. Typhi and the immune responses that might be relevant to protection against this infection are poorly understood. Unlike *S*. Typhi, non-typhoidal *Salmonella* serovar, *S*. Typhimurium, which causes only localized gastroenteritis in humans, produces a systemic disease in mice that is analogous to human typhoid. Comparative analysis of the genomes of these two closely related *Salmonella* serovars has shown that these two share close to 90% homology at the genome level. This analysis also revealed that *S*. Typhi has lost several genes due to genome degradation or formation of pseudogenes, which might have led to its host restriction [2]. How these differences contribute to host specificity and different clinical manifestations is not clear. Recent studies have shown that *S*. Typhimurium expresses a serine protease, GtgE, that degrades Rab29 and Rab32 which are known to regulate intracellular trafficking. The degradation of these Rabs promotes survival of *S*. Typhimurium inside macrophages, which led to the suggestion that these Rabs might also be involved in delivering antimicrobial molecules to the *Salmonella* - containing vacuole [3]. In fact, a recent study has identified itaconate as one of the antimicrobial molecules that is delivered by Rab32 to this vacuole [4]. GtgE is not present in *S*. Typhi and its ectopic expression increases intracellular survival of this serovar inside mouse macrophages [3, 5, 6]. Therefore, absence of this enzyme might be one of the reasons responsible for the inability of *S*. Typhi to establish infection in mice.

Several studies have reported use of immune-deficient mice reconstituted with human immune cells (humanized mice) for investigating infection with *S*. Typhi [7,8]. Unlike WT mice, infection of these mice with *S*. Typhi resulted in replication of this pathogen in various organs including spleen, liver and gall bladder reaffirming that the presence of human immune cells supports infection with this *Salmonella* serovar. However, due to heterogeneities in donor individuals and variable extents of cell engraftment, this model has found limited use in carrying out long term investigations with *S*. Typhi [7, 9]. Nevertheless, what these studies have reiterated is that the inability of *S*. Typhi to establish infection in mice is primarily due to the ability of mouse cells to prevent replication and residence of this pathogen. The mechanism of this selective prevention remains unclear.

In the present study, we analyzed early inflammatory and antibacterial responses in mice infected with *S*. Typhi and *S*. Typhimurium to understand the reasons for different outcomes of infection with these two closely related serovars. Our findings suggest that generation of antibacterial activities, both intracellular and extracellular, early on with *S*. Typhi might contribute to non-establishment of infection with this serovar in mice.

## Materials and Methods

### Mice and bacterial strains

Wild type, T and B cell deficient, MyD88^-/-^ (on C57BL/6 background), CBA/J, C3H/OuJ and C3H/HeJ mice were procured from the Jackson laboratory, USA and maintained at the Small Animal Facility of the National Institute of Immunology. The experiments with mice were carried out according to the guidelines provided by the Institutional Animal Ethics Committee (IAEC). Vi negative *Salmonella* Typhi was provided by Prof. Geeta Mehta, Department of Microbiology, Lady Hardinge Medical College, New Delhi. *Salmonella* Typhimurium SL1344 strain was obtained from Prof. Emmanuelle Charpentier, Department of Microbiology and Genetics, University of Vienna, Austria (now at the Max Planck Institute for Infection Biology in Berlin). Bacteria were cultured in LB medium at 37°C with shaking at 220 rpm for 10-12h. Bacterial suspension was centrifuged at 8000xg for 5 min to pellet down bacteria. Bacterial pellet was washed two times with serum-free RPMI-1640, resuspended in RPMI-1640 and then used for mouse infection. Bacterial loads were determined by plating tissue lysates on *Salmonella*– *Shigella* (SS) agar plates. Other bacteria used in this study were obtained from the laboratories of Dr. Kanwaljit Kaur and Dr. Vinay Nandicoori, National Institute of Immunology, New Delhi. These bacteria were grown using standard culture protocols.

### Serum ELISA

Mice were infected with 5×10^6^ CFU of *S*. Typhi or *S*. Typhimurium. At 1 h, 4 h and 8 h post infection, serum samples were collected and analyzed for IL-6 (BD OptEIA™), IL-12p40 (BD OptEIA™), KC (R&D Systems) and TNF-α (BD OptEIA™) by ELISA as per manufacturer’s instructions.

### Flow cytometric analysis of cell populations in the peritoneum

Macrophages, B cells, and T cell populations in the peritoneum of uninfected, *Salmonella* Typhi and *Salmonella* Typhimurium-infected mice were analyzed by flow cytometry using cell type specific antibodies. Briefly, Peritoneal cells were adjusted to 1×10^6^ cells and stained with specific antibodies against cell surface markers CD11b (M1/70 clone), Gr1 (RB6-8C5 clone), F4/80 (BM8 clone), CD4 (GK1.5 clone), CD8 (53-6.7 clone) and B220 (RA3-6B2 clone), or isotype matched antibodies (BD, San Jose, CA) for 60 min at 4°C. After washing, 20,000 cells were analyzed in a flow cytometer (FACS Verse or Accuri C6; BD Pharminogen, USA). Single colour control stained cells were used for compensation.

### Analysis of antibacterial activity in the cell - free peritoneal lavages of infected mice

Mice were infected intraperitoneally with 5×10^6^ CFU of *S*. Typhi or *S*. Typhimurium. Uninfected mice received equivalent volume of RPMI-1640. At 4 h post injection, peritoneal lavages were collected, centrifuged at 800xg for 5 min and the supernatants were collected and passed through 0.22 µ membrane filter. 10,000 CFU of *S*. Typhi or *S*. Typhimurium (in 100 µl) were incubated with 900 µl of the cell -free peritoneal lavage in triplicate at 37°C and bacterial growth was monitored by measuring absorbance at 630 nm. In some experiments mice were treated with PMSF (1 mg/mouse) half an hour before infecting with *Salmonella*. DMSO was used as vehicle control.

### Analysis of antibacterial activity from *in vitro*-infected macrophages

6-8-week-old C57BL/6 mice were euthanised by cervical dislocation and peritoneal cells were recovered with the help of ice cold RPMI. Cells were centrifuged at 800xg for 5 min and resuspended in RPMI containing 10% FCS. Cells were plated in a 6-well plate at a density of 4×10^6^ cells per well and incubated at 37°C in 5% CO_2_. After 48 h, cells were washed 3 times with RPMI and infected with *S*. Typhi or *S*. Typhimurium at MOI 5 for 150 min. For experiments with antibiotic-treated bacteria, *S*. Typhi and *S*. Typhimurium were incubated with gentamicin (1 mg/10^8^ bacteria/ml) for 1 hour, washed to get rid of the antibiotic and then used for stimulating peritoneal cells. In some other experiments, cells were treated with PMSF (100 µM) or glycine (10 mM) for half an hour before infecting with *Salmonella*. The supernatants from uninfected cells and infected cultures were collected and passed through 0.22 µ membrane to get rid of bacteria or cells. 10,000 CFU of *S*. Typhimurium were incubated with 900 µl of these supernatants in triplicate at 37°C for 9 h and bacterial growth was determined by measuring absorbance at 630 nm.

### Analysis of LDH-release from infected macrophages

Peritoneal macrophages were infected with *Salmonella* in triplicate (5 MOI for 2.5 h). The culture supernatants were collected and analysed for lactate dehydrogenase (LDH) by a commercially available CytoTox 96 Non-radioactive Cytotoxicity Assay kit according to the manufacturer’s (Promega) instructions. Percent cell death was calculated as = (LDH released upon infection/LDH released by TritonX-100 treated cells) x 100. LDH obtained from cells upon lysis with 1% TritonX-100 was taken as 100% cell death.

### mRNA quantification

*In vitro* or *in vivo*-infected peritoneal cells were lysed using RLT-buffer (Qiagen, Hilden, Germany). Total RNA was extracted following the manufacturer’s instructions. RNA was eluted in RNase-free water (Qiagen). The quantity and quality of mRNA were checked by nanodrop. cDNA was synthesized using Thermo Script RT-PCR kit according to the instructions provided by the manufacturer (Invitrogen). Real-time PCR reactions were set up with a SYBR green power up kit (Applied Biosystems) on an Applied Biosystems 7500 Real Time Quantitative PCR System workstation. Target genes were normalised to stably expressed reference gene 18S rRNA. The mean threshold cycles were used for further analysis. For the *in vitro* study, RNA from uninfected cells was used as calibrator sample. For *in vivo* investigation, since there was a significant change in cellularity following infection with both serovars, comparative analysis was carried out between samples harvested at two different time points from *S*. Typhi - infected mice and those from *S*. Typhimurium - infected mice. PCR was performed with following sets of primers designed using NCBI/Primer-BLAST.

CRAMP: Forward 5’–CCCAAGTCTGTGAGGTTCCG Reverse 5’–CTTGAACCGAAAGGGCTGTG

Lipocalin: Forward 5’–AATGCGGTCCAGAAAAAAACA Reverse 5’–TGACCAGGATGGAGGTGACA

Feroportin: Forward 5’–TCCGTGAACTTGAATGTGAACAA Reverse 5’–GGAAGGGCTCTGCCATCTG

Hepcidin: Forward 5’–CCACCTATCTCCATCAACAGATGA Reverse 5’–TTCTTCCCCGTGCAAAGG

18S rRNA Forward 5’-CGAAAGCATTTGCCAAGAAT-3′ Reverse 5’-AGTCGGCATCGTTTATGGTC-3′

### Proteinase K digestion

Cell-free peritoneal lavage from *S*. Typhi-infected mice was treated with Proteinase K (100 µg/ml) for 1h and passed through 0.22 µ membrane. Antibacterial activity was determined by incubating 10,000 *S*. Typhimurium at 37°C with undigested and Proteinase K-digested cell-free peritoneal lavage.

### 2D gel electrophoresis of the cell-free peritoneal lavages

Mice were infected intraperitoneally with 5×10^6^ CFU of *S*. Typhi or *S*. Typhimurium. At 8h post infection, peritoneal lavages were collected and passed through 0.22µ membrane. Cell-free lavages were precipitated with trichloroacetic acid. 2D sample buffer was added to the precipitate and iso-electric focusing was carried out according to the protocol described by Adams and Gallagher [10], using GE IPGphor device. After iso-electric focusing, the strip was placed on top of SDS-polyacrylamide gel and subjected to electrophoresis in the second dimension at 40mA for 4h. The spots in the 2D gel were visualized by silver staining as described by Sasse and Gallagher [11].

### Mass Spectrometric analysis

Silver-stained 2D spot was processed for mass spectrometric analysis as described by Shevchenko et al., 2006 [12]. The protein sample was analysed using EASY-nLC system (Thermo Fisher Scientific, USA) coupled to LTQ Orbitrap-Velos mass spectrometer (Thermo Fisher Scientific). The resolution of peptide mixture was carried out using a 10 cm PicoFrit Self-Pack microcapillary column (inner diameter = 75 µm, outer dimeter = 360 µm, 15 µm tip) filled with 5 µm C18-resin (New Objective, USA). Analysis of protein sample was performed at the Mass Spectrometry Facility of the National Institute of Immunology, New Delhi.

### Statistical analysis

Statistical analyses were performed using the Graph Pad Prism 8 software package. Statistical differences were evaluated using Mann-Whitney test (for comparing two groups) or Kruskal-Wallis test (for three or more groups) followed by Dunn’s multiple comparison test for *in vivo* experiments. For *in vitro* experiments, Student’s two tailed t-test was used for comparing two groups and one-way ANOVA was used when comparing three or more than three groups, using appropriate post hoc analyses. A p value of less than 0.05 (*p < 0.05, **p < 0.01, and ***p < 0.001) was considered statistically significant. p values for the data shown in each panel have been included in figure legends. Error bars represent standard error mean.

## RESULTS

### Cell-free fluids from peritoneal cells infected with *S*. Typhi *in vivo* or *in vitro* inhibit bacterial replication

To investigate early host responses upon infection with *S*. Typhi and *S*. Typhimurium, we infected mice intraperitoneally with these two *Salmonella* serovars and analyzed peritoneal bacterial load, peritoneal cellularity and circulating cytokines. As expected, bacterial loads, both intracellular (peritoneal cells) and extracellular (cell-free peritoneal lavages), were higher in mice infected with *S*. Typhimurium (Fig. 1 A). The difference was close to 100 folds in extracellular bacteria and more than 10 folds in intracellular bacteria. Infection with *Salmonella* resulted in changes in the proportions of various cell types in the peritoneum but these differences were comparable in the two serovars. The numbers of F4/80^+^ macrophages decreased significantly most likely due to pyroptotic cell death (Fig. 1 B; 13). The numbers of CD11b^lo^Gr1^hi^ cells were also reduced with time (Fig. 1 B). On the other hand, CD11b^hi^Gr1^int^ cell proportions increased with time (Fig. 1 B). There was also reduction in the proportions B220^+^ B cells, CD4^+^ T cells and CD8^+^ T in the peritoneal cavities of mice infected with either serovar (Supplementary Fig. 1). Similar changes in cellularity have been previously reported in the peritoneal cavities and spleens of mice infected with pathogenic *Salmonella* and these have been associated with immunomodulatory roles in the establishment of infection with this pathogen [14–16].

**Figure 1:**
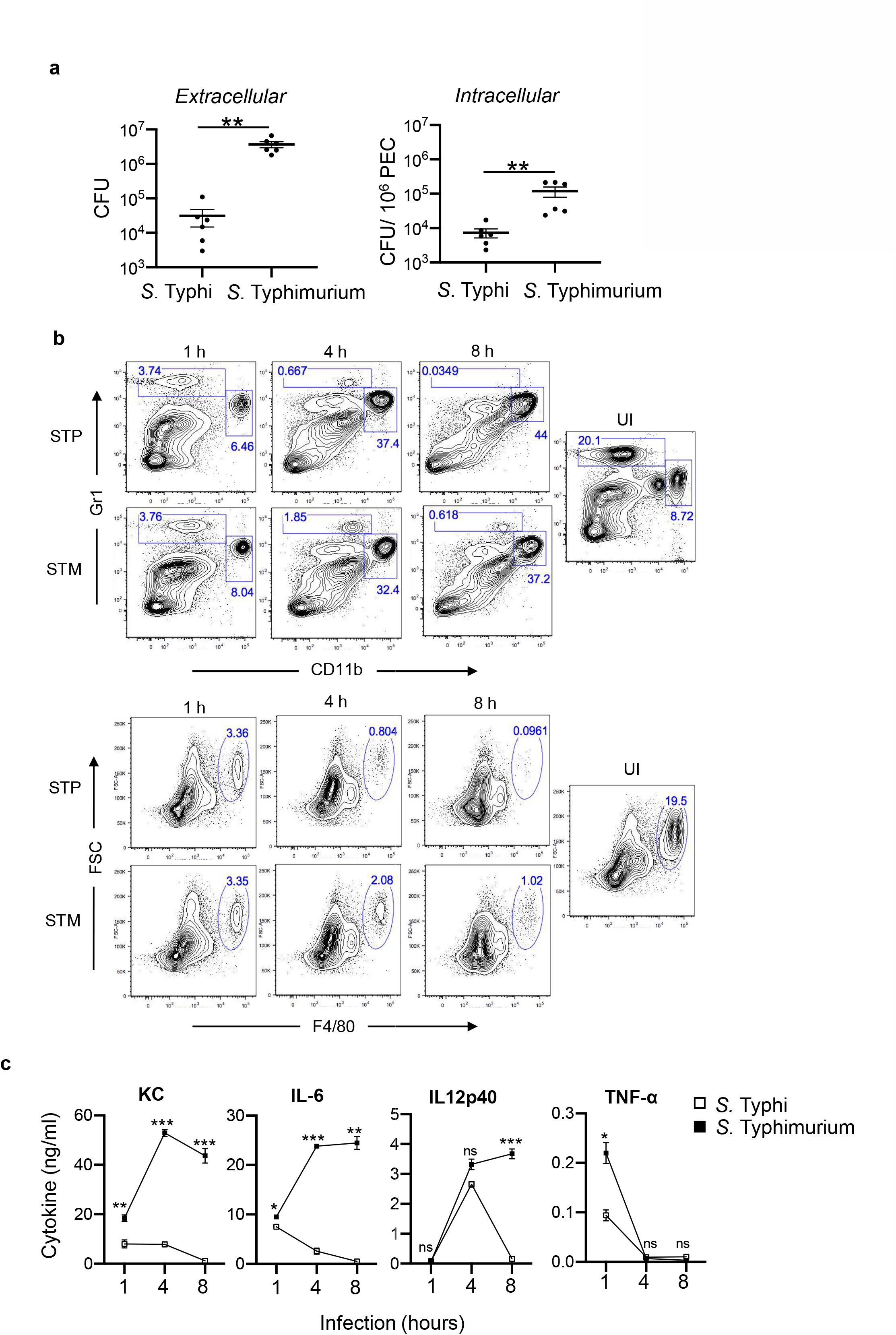
*S*. Typhi is cleared from the peritoneal cavity of mice. (**A**) C57BL/6 mice (6 per group) were infected intraperitoneally with 5×10^6^ *S*. Typhimurium or *S*. Typhi. The number of extracellular and intracellular bacteria recovered from the peritoneum 4 h after infection was determined by plating on *SS* agar plates. (**B**) C57BL/6 mice were infected intraperitoneally with 5×10^6^ *S*. Typhimurium or *S*. Typhi. Cells were isolated at 1 h, 4 h and 8 h after infection and stained with fluorophore-labelled antibodies against CD11b (FITC-labelled), Gr1 (PE-labelled) and F4/80 (V450-labelled). 20,000 cells were acquired in a flow cytometer (BD Verse). The results obtained with three mice in each group are representative of two independent experiments (UI - Uninfected, STP - *S*. Typhi, STM - *S*. Typhimurium). (**C**) Sera were collected at 1 h, 4 h and 8 h after intraperitoneal infection with 5×10^6^ of *S*. Typhimurium or *S*. Typhi (4 mice per group), and the levels of different cytokines were determined by ELISA. Data representative of 2-3 independent experiments is shown as mean ± sem. * p<0.05, ** p< 0.01, *** p<0.005, ns – not significant.

In contrast to cellularity, there were significant differences in the levels of cytokines upon infection of mice with these two *Salmonella* serovars. *S*. Typhimurium-infected mice showed significantly higher levels of IL-6, IL-12p40 and KC, as compared to *S*. Typhi-infected mice (Fig. 1 C). IL-6 was comparable at 1h post infection in mice infected with either serovar (Fig. 1 C); the levels increased further at 4h and 8h in *S*. Typhimurium-infected mice but dropped to baseline in *S*. Typhi-infected mice (Fig. 1 C). In a similar fashion, IL-12p40 levels were comparable at 1h and 4h of infection, increased further at 8h in *S*. Typhimurium–infected mice while reducing close to baseline in *S*. Typhi-infected mice (Fig. 1 C). KC levels were 5 and 30 fold higher at 4h and 8h respectively in *S*. Typhimurium–infected mice (Fig. 1 C). Serum TNF-α at 1h post infection was 2-fold higher in *S*. Typhimurium–infected mice as compared to *S*. Typhi-infected mice (Fig. 1 C). While some of these cytokines are involved in clearance of bacteria early on, their increased levels with the progression of infection serve as an indicator of increased bacterial load and inflammation [17].

The reduction in extracellular as well as intracellular bacterial load in mice infected with *S*. Typhi suggested that infection with this serovar might induce antibacterial activities in the peritoneum. To explore this possibility, we first determined antibacterial activity in cell-free peritoneal lavages prepared from infected mice. This analysis showed striking differences between peritoneal fluids from *S*. Typhi-infected mice and *S*. Typhimurium-infected mice (Fig. 2 A). Peritoneal fluids from *S*. Typhi-infected mice prevented growth of *Salmonella* (*S*. Typhi as well as *S*. Typhimurium) significantly better than those obtained from *S*. Typhimurium-infected mice (Fig. 2 A). The production of this activity was not dependent on TLR activation as peritoneal fluids obtained from MyD88-deficient mice infected with *S*. Typhi also showed antibacterial activity (Supplementary Fig. 2). T and B lymphocytes also did not contribute to generation of this antibacterial factor(s) as inhibition in bacterial growth was also observed with cell-free peritoneal lavages obtained from *S*. Typhi-infected mice deficient in either of these two cell types (Supplementary Fig. 3). Significantly, the ability of this antibacterial activity to prevent bacterial growth was not restricted to *Salmonella*; this activity suppressed replication of other Gram-negative bacteria including *Escherichia coli, Klebsiella pneumoniae* and *Pseudomonas aeruginosa*, Gram-positive bacteria such as *Staphylococcus aureus* and *Listeria monocytogenes* as well as acid fast bacillus, *Mycobacterium smegmatis* (Supplementary Fig. 4). The ability to inhibit *Salmonella* growth was also seen with peritoneal fluids from mice infected with Vi positive *S*. Typhi (data not shown).

**Figure 2:**
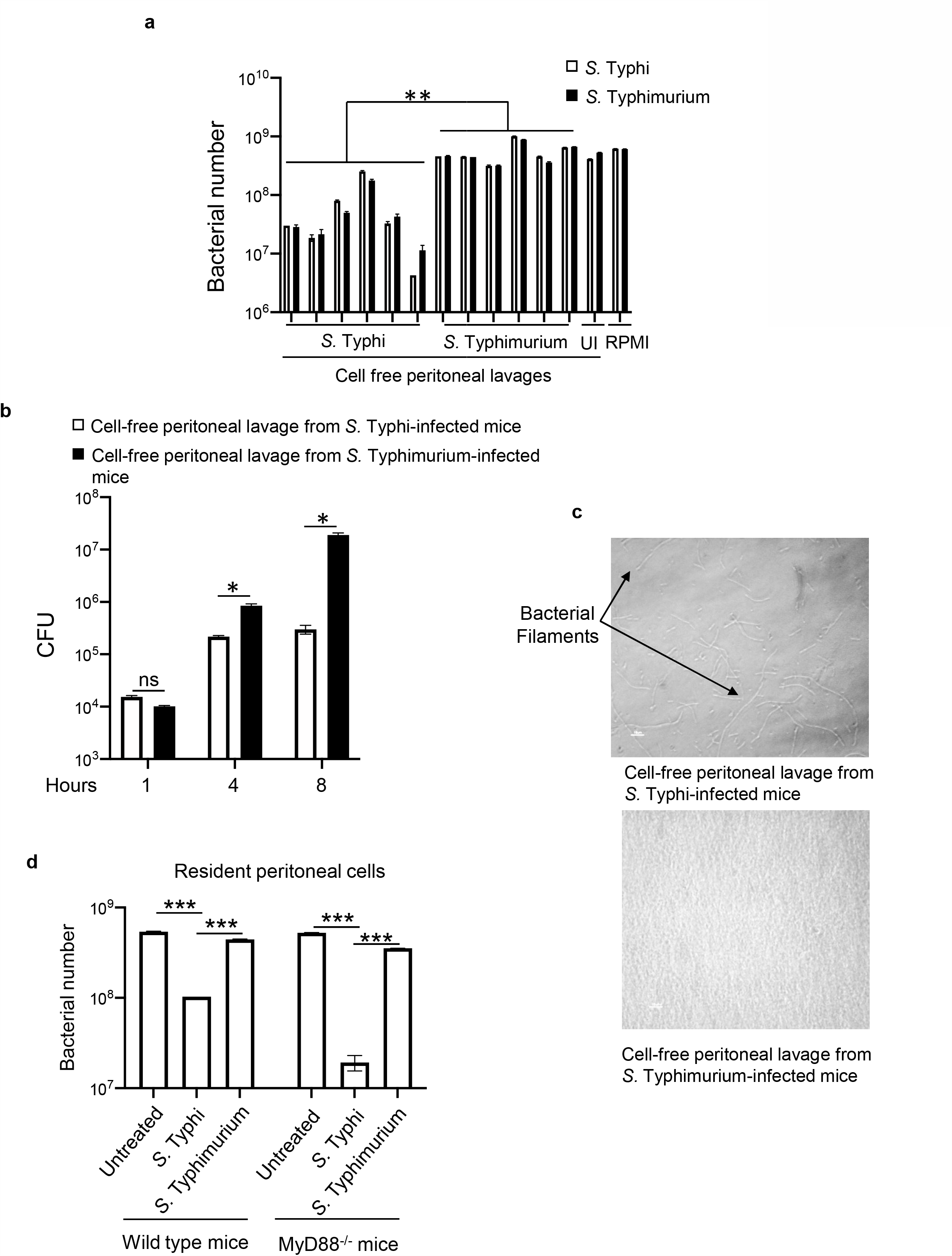
Cell-free peritoneal fluids from mice infected with *S*. Typhi and cell-free supernatants from macrophages infected *in vitro* with *S*. Typhi inhibit bacterial growth *ex vivo*. (**A**) WT C57BL/6 mice were infected intraperitoneally with 5×10^6^ CFU of *S*. Typhi or *S*. Typhimurium. Peritoneal lavages were collected from uninfected and infected mice (4 h after infection) and filtered through 0.22 µ membrane. 10,000 *S*. Typhi or *S*. Typhimurium were grown in triplicate for 23 h at 37°C in RPMI or in cell-free peritoneal lavages derived from uninfected and infected mice. Bacterial growth was determined by measuring absorbance at 630 nm. Each bar in this figure represents one mouse (UI – uninfected). (**B**) *S*. Typhimurium were incubated with cell-free peritoneal lavage from *S*. Typhi or *S*. Typhimurium-infected mice (3 mice per group) for different time points (1 h, 4 h and 8 h) and bacterial numbers were determined by plating on SS agar plates. (**C**) 10,000 *S*. Typhimurium were incubated in triplicate with peritoneal lavage from *S*. Typhi or *S*. Typhimurium-infected mice at 37°C. After 21 h, bacteria were observed under a microscope (Nikon TE2000) and images were acquired using ACT software. The results are from two independent experiments. (**D**) Peritoneal cells pooled from 3 WT or MyD88^-/-^ C57BL/6 mice were seeded in a 6 well plate (4×10^6^ per well). 48 h later, cells were infected in triplicate with *S*. Typhi or *S*. Typhimurium. Cell supernatants were collected at 4 h after infection and filtered through a 0.22 µ membrane. 10,000 *S*. Typhimurium were incubated with these supernatants at 37°C. Bacterial growth was determined after 9 h by measuring absorbance at 630 nm. Data representative of 2 independent experiments is shown as mean ± sem. * p value <0.05, ** p value <0.01, *** p<0.005, ns – not significant.

Antibacterial activity obtained from *S*. Typhi-infected mice did not readily inhibit bacterial replication in the first one hour of incubation but by 4 h, the difference in antibacterial activity between peritoneal fluids obtained from *S*. Typhi and *S*. Typhimurium-infected mice was very evident and it increased further by 8 h (Fig. 2 B). Bacterial numbers in *S*. Typhi – derived lavage at 8 h post incubation were close to 50 times lower as compared to those in *S*. Typhimurium – derived lavage. However, bacterial numbers neither reduced nor increased in presence of the cell-free lavage from *S*. Typhi – infected mice between 4 h and 8 h suggesting that this antibacterial activity may be largely bacteriostatic although simultaneous bactericidal effects can’t be ruled out. Importantly, the antibacterial activity obtained from *S*. Typhi – infected mice was also effective against *S*. Typhimurium isolated from infected mice suggesting that unlike resistance to other antimicrobial peptides such as polymyxin B, susceptibility to this activity is not altered in the course of infection (Supplementary Fig. 5) [18]. These results established that infection of mice with *S*. Typhi is accompanied by generation of extracellular antibacterial factor(s), and suggested that the innate immune cells and not the lymphocytes were the likely source of these factors. Microscopic examination revealed filamentation of *S*. Typhimurium in presence of antibacterial factor(s) present in the peritoneal fluids of *S*. Typhi-infected mice (Fig. 2 C). Interestingly, *S*. Typhi did not undergo filamentation upon treatment with this antibacterial factor(s) even though its growth was inhibited as much as that of *S*. Typhimurium (data not shown; Fig. 2 A), the reasons for which are not clear.

The induction of antibacterial activity with *S*. Typhi was also observed during its *in vitro* infection of peritoneal macrophages. Cell-free supernatants prepared from *S*. Typhi-infected cells showed higher antibacterial activity as compared to those obtained from cells infected with *S*. Typhimurium (Fig. 2 D). This induction required metabolically active bacteria as cells incubated with gentamycin-treated *S*. Typhi did not release any detectable antibacterial activity (Supplementary Fig. 6). Similar to what was observed during *in vivo* infection, the production of this antibacterial activity was not dependent on the activation of MyD88-dependent TLR signaling (Fig. 2 D). Moreover, the induction of antibacterial activity was not restricted to C57BL/6 mice; it was also observed during *S*. Typhi - infection of macrophages from other strains of mice including CBA/J, C3H/OuJ and LPS-hyporesponsive strain, C3H/HeJ (Supplementary Fig. 7).

### Induction of extracellular antibacterial activity with *S*. Typhi is dependent on serine protease

Serine proteases produced by macrophages and neutrophils have been previously shown to be involved in generating active antimicrobial peptides [19–25]. Therefore, we examined the role of these proteases in regulating production of antibacterial activity with *S*. Typhi. Treatment of mice with serine protease inhibitor, PMSF, prior to infection significantly reduced the ability of peritoneal fluid from *S*. Typhi-infected mice to inhibit replication of *Salmonella ex vivo* (Fig. 3 A). Similar reduction in this activity was seen during infection of peritoneal cells with *S*. Typhi *in vitro* (Fig. 3 B). These results indicated that one or more serine proteases might be contributing to the production of antibacterial activity by cells infected with *S*. Typhi. At concentrations used in this study, PMSF did not show any direct effect on bacterial growth (data not shown).

As infection with *Salmonella* is known to bring about pyroptotic cell death that is mainly driven by caspase-1 [13], we also looked at possible role of this cell death in the release of this antibacterial activity. *S*. Typhi did not differ from *S*. Typhimurium in bringing about cell death of macrophages (Supplementary Fig. 8). Interestingly, treatment of cells with glycine, which prevents cell lysis by blocking plasma membrane permeabilization [26], significantly reduced antibacterial activity in cell-free supernatants from *S*. Typhi-infected cells showing that cell death was required for the release of this activity (Fig. 3 C, D). PMSF, which inhibited generation of antibacterial activity during *in vivo* as well as *in vitro* infection (Fig. 3 A, B), did not affect cell death produced by *S*. Typhi (Supplementary Fig. 9). These results suggested that the production of antibacterial factor(s) by peritoneal cells upon infection with *S*. Typhi is regulated by one or more serine proteases, and its release from cells requires cell death. Alternatively, these proteases might be generating this activity extracellularly after getting released as a result of cell death.

**Figure 3:**
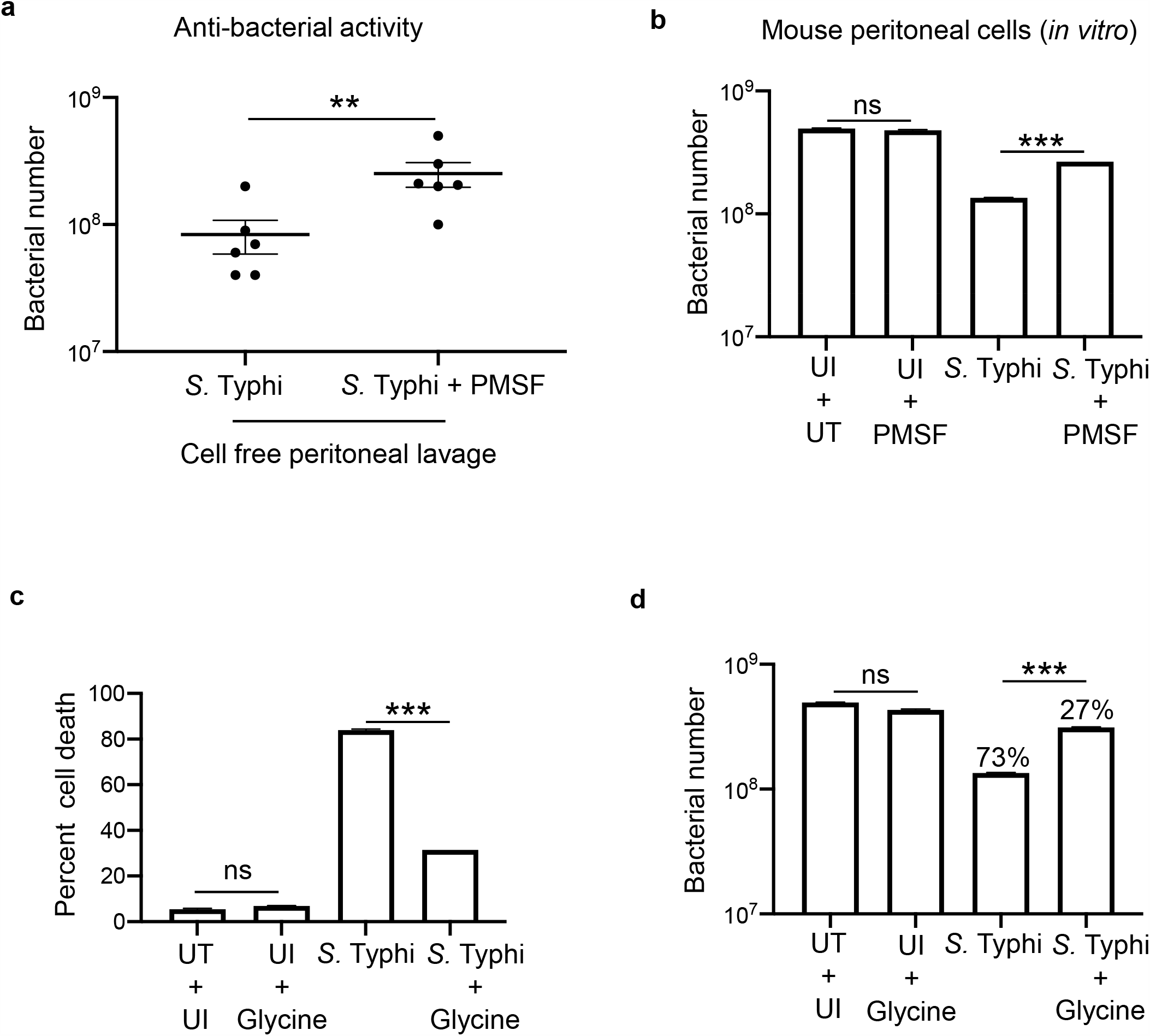
Induction of antibacterial factor(s) with *S*. Typhi is dependent on serine protease activity, and its release is dependent on cell death. (**A**) C57BL/6 mice were pretreated with serine protease inhibitor, PMSF, (1 mg/mouse) or with the vehicle (DMSO) and 1 h later, mice were infected with *S*. Typhi. *S*. Typhimurium was incubated for 21 h with cell-free peritoneal lavages (obtained after 4 hours of infection) from untreated and PMSF-pretreated, *S*. Typhi-infected mice. Bacterial growth was determined by measuring absorbance at 630 nm. (**B**) Peritoneal macrophages were pretreated with PMSF (100 µM) or with the vehicle (DMSO) in triplicate, and 30 min later, cells were infected with *S*. Typhi for 2.5 h. Cell-free supernatants were collected and passed through 0.22 µ membrane. 10,000 *S*. Typhimurium were incubated in triplicate with these cell-free supernatants and bacterial growth was determined by measuring absorbance at 630 nm. The results are representative of two independent experiments. (UI **–** uninfected, UT **–** untreated). (**C, D**) Resident peritoneal cells from C57BL/6 were seeded in a 6 well plate (4×10^6^ per well). 48 h later, cells were pretreated with glycine (10 mM) in triplicate and 30 min later, these were infected with *S*. Typhi for 2.5h. Cell death was determined by release of LDH in the supernatant. Culture supernatants collected at 2.5h were passed through 0.22 µ membrane and evaluated for antibacterial activity by incubating with *S*. Typhimurium. Bacterial growth was determined after 9 h by measuring absorbance at 630 nm. The results are from two independent experiments. Data shown is represented as mean ± sem (UI – uninfected, UT– untreated). ** p value <0.01, *** p<0.005, ns – not significant.

Preliminary characterization of the antibacterial activity in the peritoneal fluids of *S*. Typhi-infected mice showed that it was more than 10 KDa and proteinaceous in nature as it lost its activity upon digestion with Proteinase K (Supplementary Fig. 10 a, b). Analysis by 2D gel electrophoresis showed that a large majority of proteins were conserved in the peritoneal fluids obtained from mice infected with *S*. Typhi and *S*. Typhimurium (Supplementary Fig. 10 c). A very prominent 50 KDa spot present only in the peritoneal fluid of *S*. Typhi-infected mice was identified as cleaved serotransferrin by mass spectrometry (Supplementary Fig. 10 c).

### Infection with *S*. Typhi results in increased expression of lipocalin and ferroportin mRNAs

To analyze possible intracellular antibacterial activities, we focused on molecules involved in regulating iron, which is known to play a crucial role in controlling *Salmonella* replication in macrophages [27–29]. We examined mRNA expression of lipocalin, ferroportin and hepcidin, and also analyzed expression of murine cathelin-related antimicrobial peptide (CRAMP). Lipocalin, ferroportin and hepcidin regulate the levels and metabolism of iron, and CRAMP possesses direct antibacterial activity [27–29]. Lipocalin checks bacterial growth by binding iron-loaded siderophores while ferroportin transports iron from inside of the cell to outside of the cell thus making it unavailable for intracellular bacteria during infection [27,28]. Hepcidin regulates iron levels by binding to ferroportin [30]. Considering significant changes in the proportions of various cell types including F4/80^+^ macrophages (Fig. 1 B), which might be the first cell type that gets infected with *Salmonella* during intraperitoneal infection, we didn’t determine expression of these molecules in the peritoneal cells from uninfected mice. Comparison was carried out between mice infected with *S*. Typhi and those infected with *S*. Typhimurium. The expression of lipocalin and ferroportin was similar at 4 h in peritoneal cells isolated from *S*. Typhi and *S*. Typhimurium-infected mice. However, at 8 h of infection, these levels reduced significantly in *S*. Typhimurium-infected mice but not in *S*. Typhi – infected mice (Fig. 4 A). The levels of hepcidin mRNA were not different in the two sets of infected mice (Fig. 4 A). The levels of CRAMP mRNA were similar in the two sets of mice at 4 h of infection. However, by 8 h, these levels reduced in *S*. Typhi-infected mice but increased significantly in *S*. Typhimurium-infected mice (Fig 4 A).

The expression of CRAMP, lipocalin, ferroportin and hepcidin was also analysed during *in vitro* infection of peritoneal macrophages with these two *Salmonella* serovars (Fig. 4 B). The mRNA levels of CRAMP, ferroportin, and hepcidin increased significantly in macrophages infected with either serovar (Fig. 4 B). Unlike what was seen with peritoneal cells from infected mice, the mRNA expression of CRAMP was similar in peritoneal cells infected with *S*. Typhi or *S*. Typhimurium *in vitro* (Fig. 4 B). Lipocalin-2 increased 2-3 times upon infection with *S*. Typhi but not with *S*. Typhimurium (Fig. 4 B). The levels of ferroportin and hepcidin mRNAs increased upon infection with *S*. Typhi as well *S*. Typhimurium but the increase was significantly more with *S*. Typhi (Fig. 4 B). The reasons for differences between *in vivo* and *in vitro* analyses in the pattern of CRAMP and hepcidin expression are not clear at the moment. These might be due to differences in the responses of different cell types recruited to the peritoneum during infection with the two *Salmonella* serovars. It is also possible that their expression is modulated by inflammatory cytokines, which show significant differences in mice infected with the two serovars (Fig. 1 C). Irrespective of these differences, the results showed that infection with *S*. Typhi induces higher and sustained expression of lipocalin and ferroportin, which might contribute to clearance of this serovar in mice.

**Figure 4:**
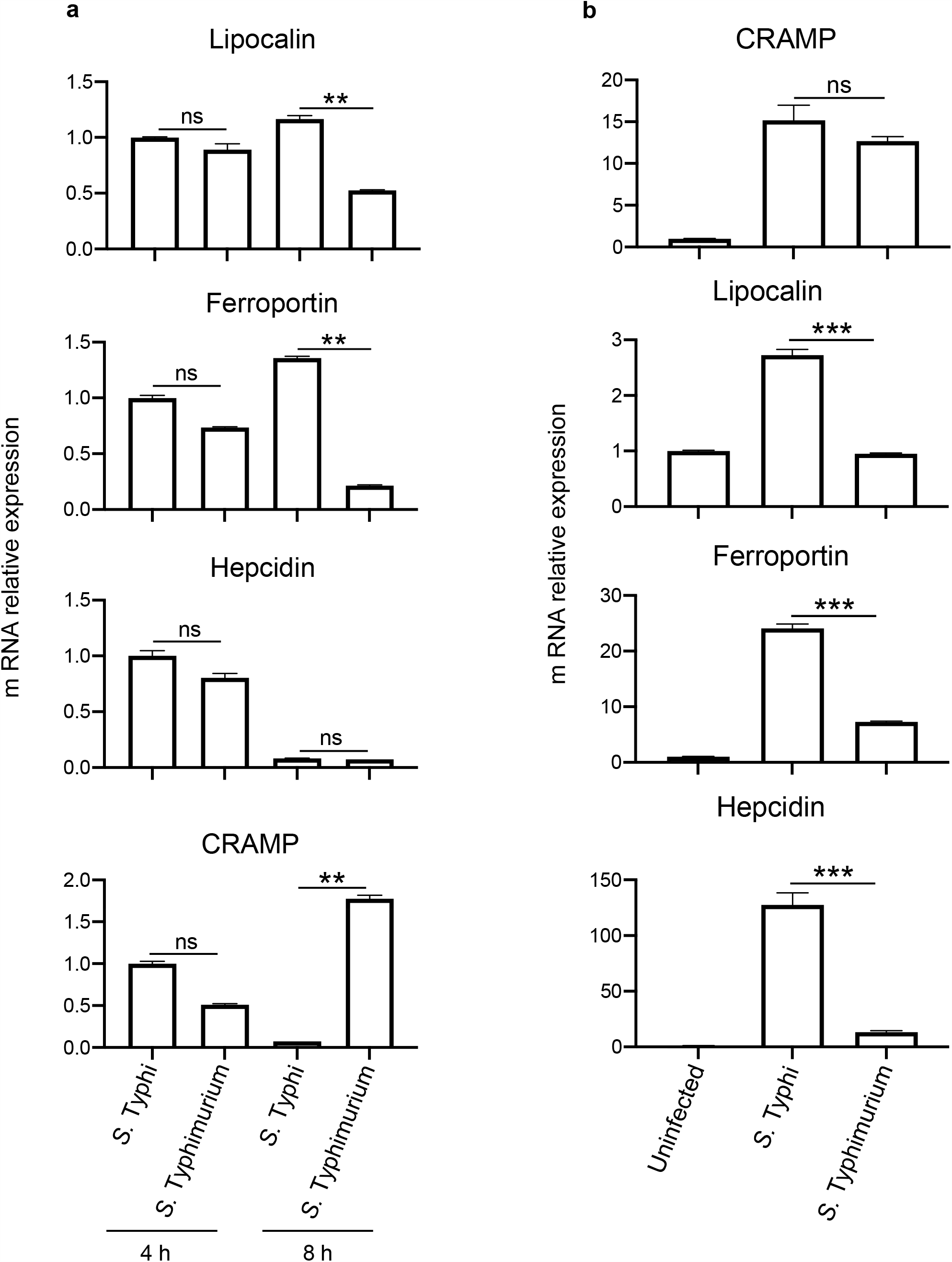
Induction of antimicrobial molecules in peritoneal cells from mice infected with *S*. Typhi or *S*. Typhimurium. (**A**) Mice were infected intraperitoneally with 5×10^6^ CFU of *S*. Typhi or *S*. Typhimurium. RNA was isolated from peritoneal cells collected after 4 h and 8 h of infection. mRNA levels of Lipocalin, Ferroportin, Hepcidin and CRAMP, were determined using real time PCR. Data are representative of three mice per group from two independent experiments. (**B**) Mouse resident peritoneal macrophages were infected *in vitro* in triplicate with 10 MOI of *S*. Typhi or *S*. Typhimurium for 1 h. Cells were washed to remove extracellular bacteria and incubated for another 24h in presence of gentamycin (100 µg/ml). CRAMP, Lipocalin, Ferroportin and Hepcidin mRNA levels were determined using real time PCR. Data representative of two independent experiments is shown as mean ± sem. ** p value <0.01, *** p<0.005, ns – not significant.

## DISCUSSION

Our results reveal induction of extracellular and intracellular antibacterial activities during clearance of *S*. Typhi from the peritoneal cavity of mice. The production of extracellular pan anti-bacterial activity seems to be in part driven by one or more serine proteases. The exact identity of the factor(s) present in this activity is not established at the moment. Preliminary analysis showed presence of cleaved serotransferrin in this activity. This molecule could contribute to inhibiting bacterial growth through depletion of iron as its proteolytically cleaved high and low molecular weight fragments have been previously shown to have antibacterial and proinflammatory activities respectively [31,32]. Significantly, a serine protease-generated antibacterial activity similar to the one found during *in vivo* infection was also observed during *in vitro* infection of macrophages with *S*. Typhi. Serine proteases have been shown to participate either directly or through generation of antimicrobial molecules in host defense against bacterial pathogens [25]. Interestingly, the release of the antibacterial activity from *S*. Typhi-infected macrophages was found to be dependent on pathogen-induced cell death. Considering that the magnitudes of macrophage cell death produced by *S*. Typhi and *S*. Typhimurium were not different, this observation raises an interesting possibility that cell death produced by the two *Salmonella* serovars might be qualitatively different.

Intracellularly, both *in vivo* and *in vitro*, infection with *S*. Typhi resulted in increased induction of lipocalin and ferroportin mRNAs. Lipocalin is expressed by macrophages, neutrophils, and epithelial cells and is up-regulated in response to infectious and inflammatory stimuli [33]. It inhibits bacterial growth by impeding siderophore-mediated iron uptake [28]. Ferroportin is the only known iron transporter which can transport iron from intracellular milieu to the extracellular environment thereby limiting intracellular iron levels [34]. The acquisition of iron is known to be crucial for survival of *Salmonella* in macrophages [35], and *S*. Typhi has been shown to be more sensitive to iron limitation than *S*. Typhimurium [36]. *In vivo* administration of iron to mice has been previously reported to promote systemic replication of *S*. Typhi [37]. In J774 murine macrophage cell line, ferroportin overexpression lead to iron deprivation and subsequent reduction in intracellular growth of *Salmonella* [27]. Therefore, it is likely that iron deprivation due to enhanced lipocalin and ferroportin expression during infection with *S*. Typhi infection selectively restricts growth of this *Salmonella* serovar in mouse cells. The reasons for lack of upregulation of these two antibacterial proteins during infection with *S*. Typhimurium need to be investigated.

Spanò and Galán have suggested that *S*. Typhi might be cleared in mouse macrophages by one or more antibacterial factors that are delivered to *Salmonella*-containing vacuole by Rab32 [6]. In a recent study from the Galan laboratory, Chen et al. have shown that one of the antibacterial molecules delivered by Rab32 to the *Salmonella*-containing molecule in macrophages is itaconate [4]. Rab29 and Rab32 are specifically cleaved by a protease, GtgE, present in *S*. Typhimurium but not in *S*. Typhi [3,6]. Ectopic expression of this enzyme in *S*. Typhi resulted in its increased replication inside mouse macrophages [6]. The study by Chen et al. suggested that Rab32 might deliver additional antibacterial factors to bring about clearance of pathogenic *Salmonella* [4]. Our results indicate that antibacterial molecules involved in the regulation of iron might also critically contribute to the inability of *S*. Typhi to establish infection in mice [6].

Taken together, our findings suggest that early anti-bacterial activities generated selectively with *S*. Typhi might play a major role in preventing establishment of infection with this *Salmonella* serovar in mice. Further investigations on the identity of extracellular antibacterial activity and the mechanism involved in the induction of lipocalin and ferroportin should facilitate development of a suitable animal model for *S*. Typhi infection.

## Notes

### Data Availability Statement

All relevant data are within the paper and its supporting information files.

## Acknowledgments

We thank Drs Rahul Pal, Devinder Sehgal, Perumal Nagarajan, Chinmay Mukhopadhyay and Naeha Subramanian for their valuable suggestions in the course of this study, and members of the Ayub laboratory for insightful discussions. We are thankful to Basanti Malakar and Shanta Sen for Mass Spectrometric analysis.

## Author contributions

A.Q. conceived and supervised the study; J.Y. and A.Q. designed experiments; J.Y. performed experiments and prepared data for publication; J.Y. and A.Q. analyzed the data, and wrote the manuscript.

