## Supplementary Figures for "Early induction of antibacterial activities distinguishes response of mice to infection with non-permissive from response to permissive *Salmonella*" (from another version of this preprint: version 1, 2020-12-07)

#### Supplementary Figure 1

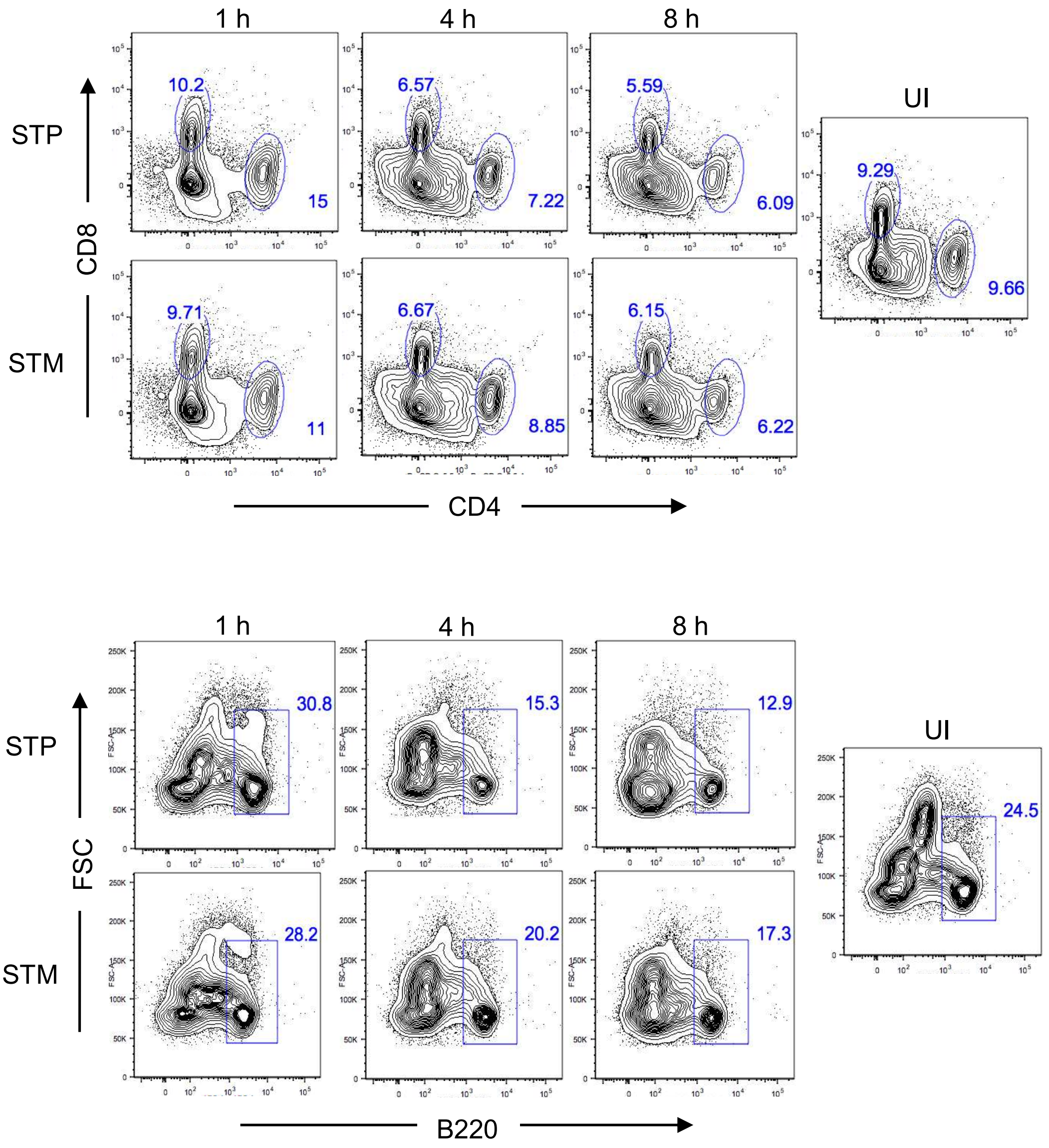


**Supplementary Fig. 1: Infection of mice with *S.* Typhi or *S.* Typhimurium results in similar changes in the proportions of peritoneal T and B lymphocytes.**
Peritoneal cells from uninfected mice and mice infected with S. Typhi or S. Typhimurium were stained with fluorophore-labeled antibodies against CD4 (perCP-cy5.5-labelled), CD8 (PE-Cy7-labelled) and B220 (APC-labelled). 20,000 cells were analyzed in a flow cytometer (BD Verse). The results obtained from 3 mice in each group are representative of two independent experiments (UI – uninfected, STP – *S.* Typhi – infected;, STM - *S.* Typhimurium – infected).

#### Supplementary Figure 2

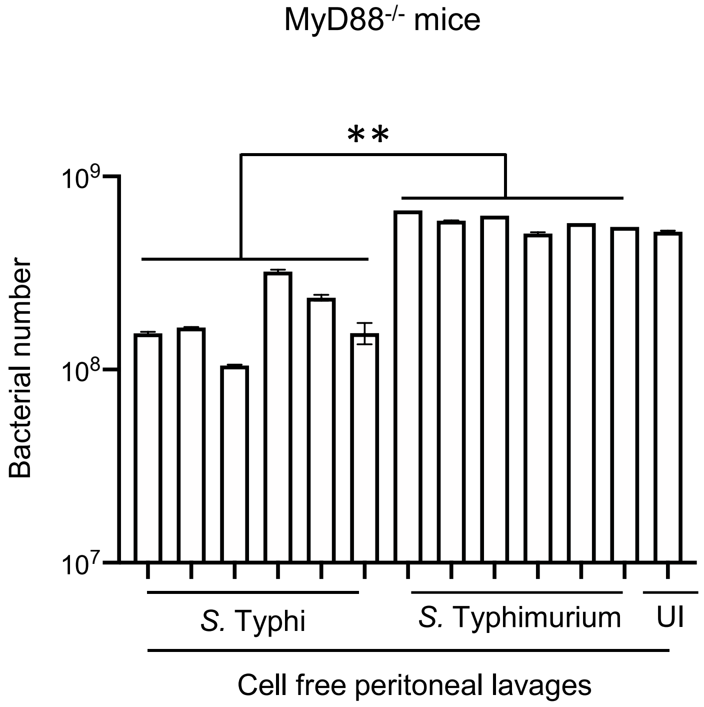


**Supplementary Fig. 2: MyD88-dependent TLR siganling does not participate in the induction of antibacterial activity during infection of mice with *S*. Typhi.**
MyD88 deficient C57Bl/6 mice were infected i.p. with $5x10^6$ *S.* Typhimurium or *S.* Typhi. Peritoneal lavages were collected at 4 h after infection. 10,000 *S.* Typhimurium were incubated at 37ºC with cell-free peritoneal lavages derived from uninfected and infected mice. Bacterial growth was measured after 22 h by determining absorbance at 630 nm. Data representative of two experiments is shown as mean ± sem. ** p value <0.01.

#### Supplementary Figure 3

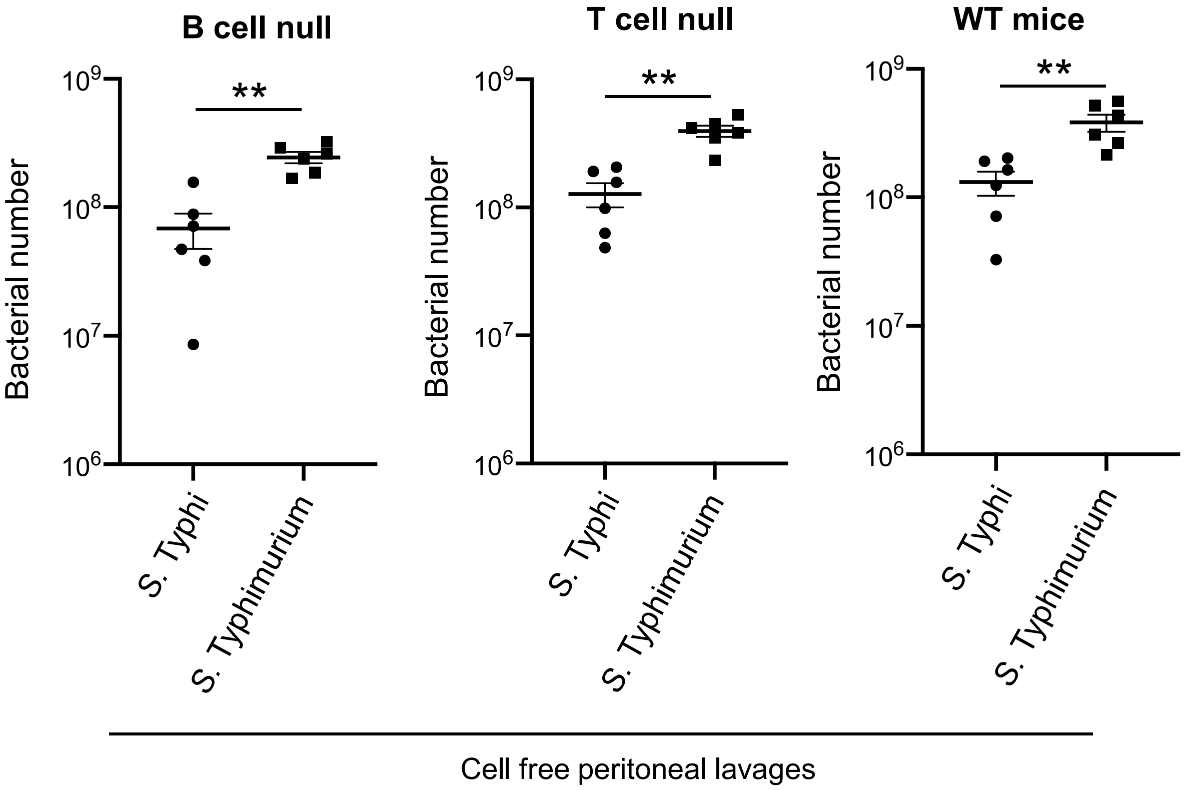


**Supplementary Fig. 3: T and B cells don't contribute to antibacterial activity generated in mice upon infection with *S*. Typhi**.

μMT$^{-/-}$, TCRβ$^{-/-}$and WT mice were infected i.p. with $5x10^6$ *S.* Typhimurium or *S.* Typhi. Peritoneal lavages were collected at 4h after infection. *S.* Typhimurium was incubated at 37°C with cell - free peritoneal lavages derived from *S*. Typhi or *S*. Typhimurium-infected mice. Bacterial growth was measured after 22 h by determining absorbance at 630 nm. Data representative of two independent experiments is shown as mean ± sem. ** p value <0.01.

### Supplementary Figure 4

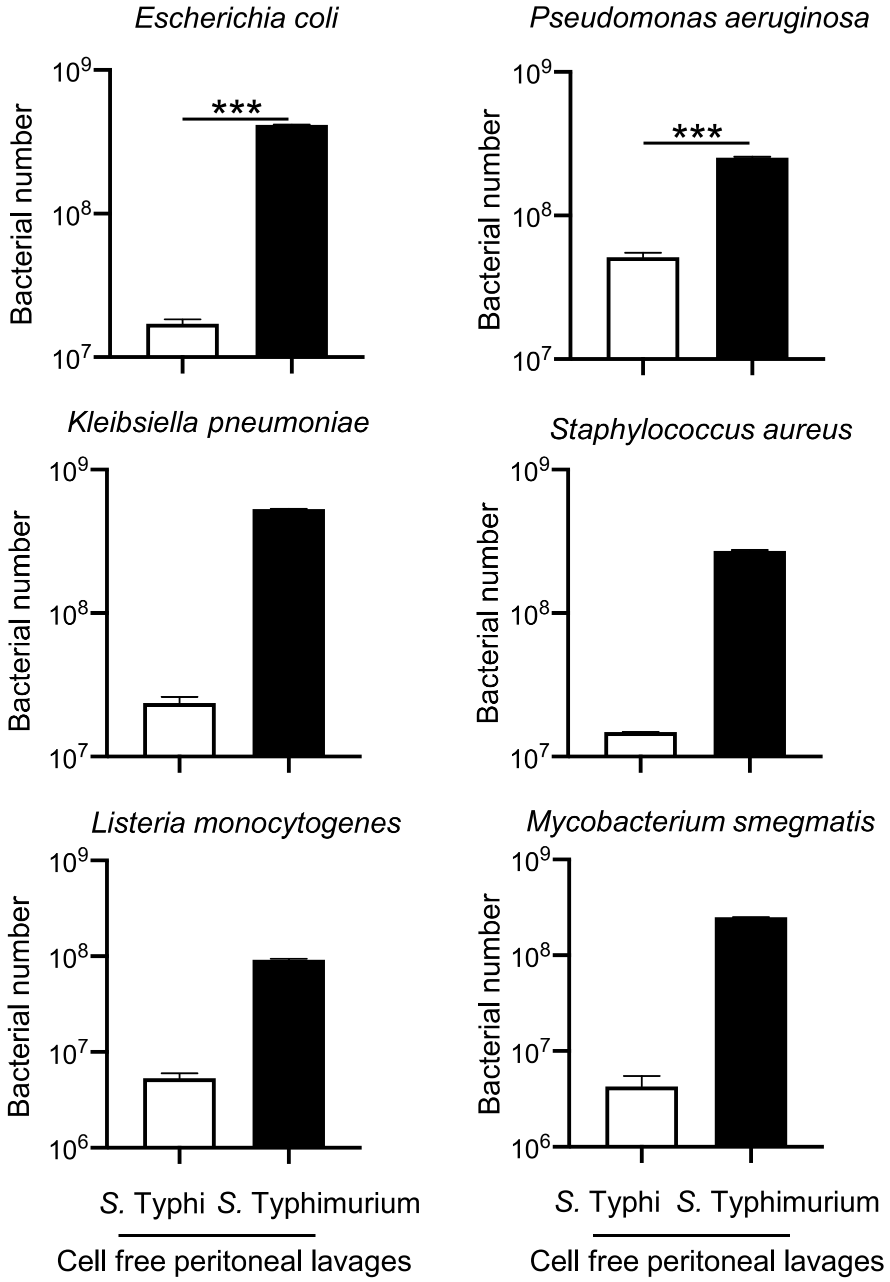


**Supplementary Figure 4: Antibacterial activity elicited in mice infected with *S.* Typhi inhibits growth of gram positive as well as gram negative bacteria.**
$10^4$ *Escherichia coli, Pseudomonas aeruginosa, Klebsiella pneumoniae, Staphylococcus aureus*, *Listeria monocytogenes or Mycobacterium smegmatis* were incubated with cell-free peritoneal lavages obtained from *S.* Typhi or *S.* Typhimurium - infected mice and bacterial growth was determined by measuring absorbance at 630 nm. Data representative of two independent experiments is shown as mean ± sem. *** p value <0.005.

#### Supplementary Figure 5

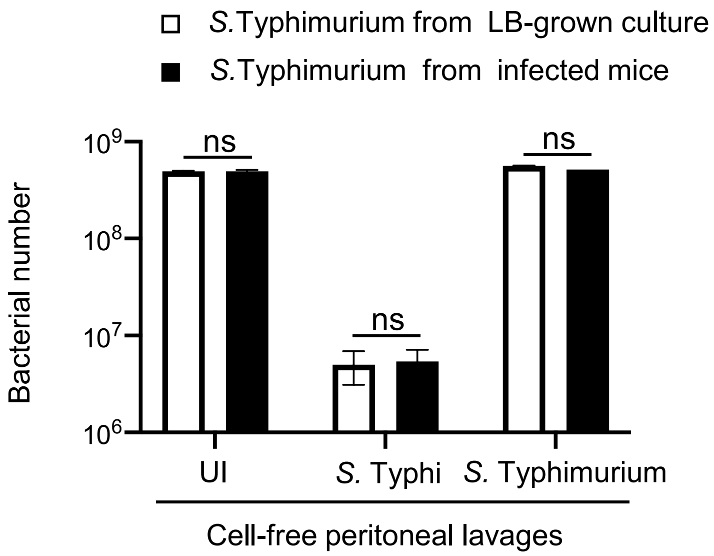


**Supplementary Figure 5: Cell free peritoneal lavage from *S.* Typhi – infected mice is also effective in inhibiting growth of *S.* Typhimurium isolated from the spleens of infected mice.**

10,000 *S.* Typhimurium obtained from LB-grown culture or from mice on day 4 of ip infection were grown for 12h at 37°C in cell-free peritoneal lavages derived from uninfected or *Salmonella*-infected mice. Bacterial growth was determined by measuring absorbance at 630 nm. Data representative of three independent experiments and shown as mean ± sem. UI – uninfected; ns – not significant.

#### Supplementary Figure 6

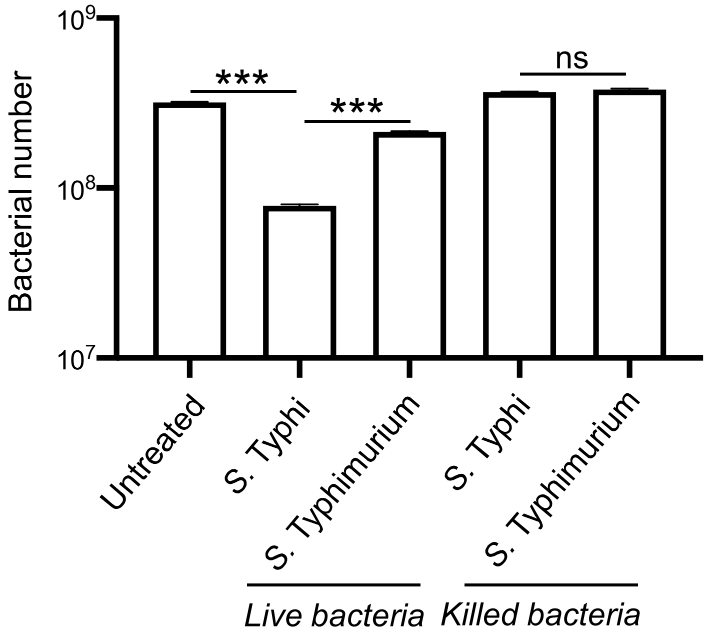


**Supplementary Fig. 6: Induction of antibacterial activity requires metabolically active bacteria.**

Resident peritoneal cells from C57BL/6 mice were seeded in a 6 well plate ($4x10^6$ per well). 48 h later, cells were infected with live or gentamicin - treated *S.* Typhi or *S.* Typhimurium. Cell supernatants were collected at 2.5 h after infection and filtered through a 0.22 µ membrane. 10,000 *S.* Typhimurium were incubated with these supernatants at 37ºC. Bacterial growth was determined after 9h by measuring absorbance at 630 nm. Data representative of two independent experiments is shown as mean ± sem. *** p value <0.05.

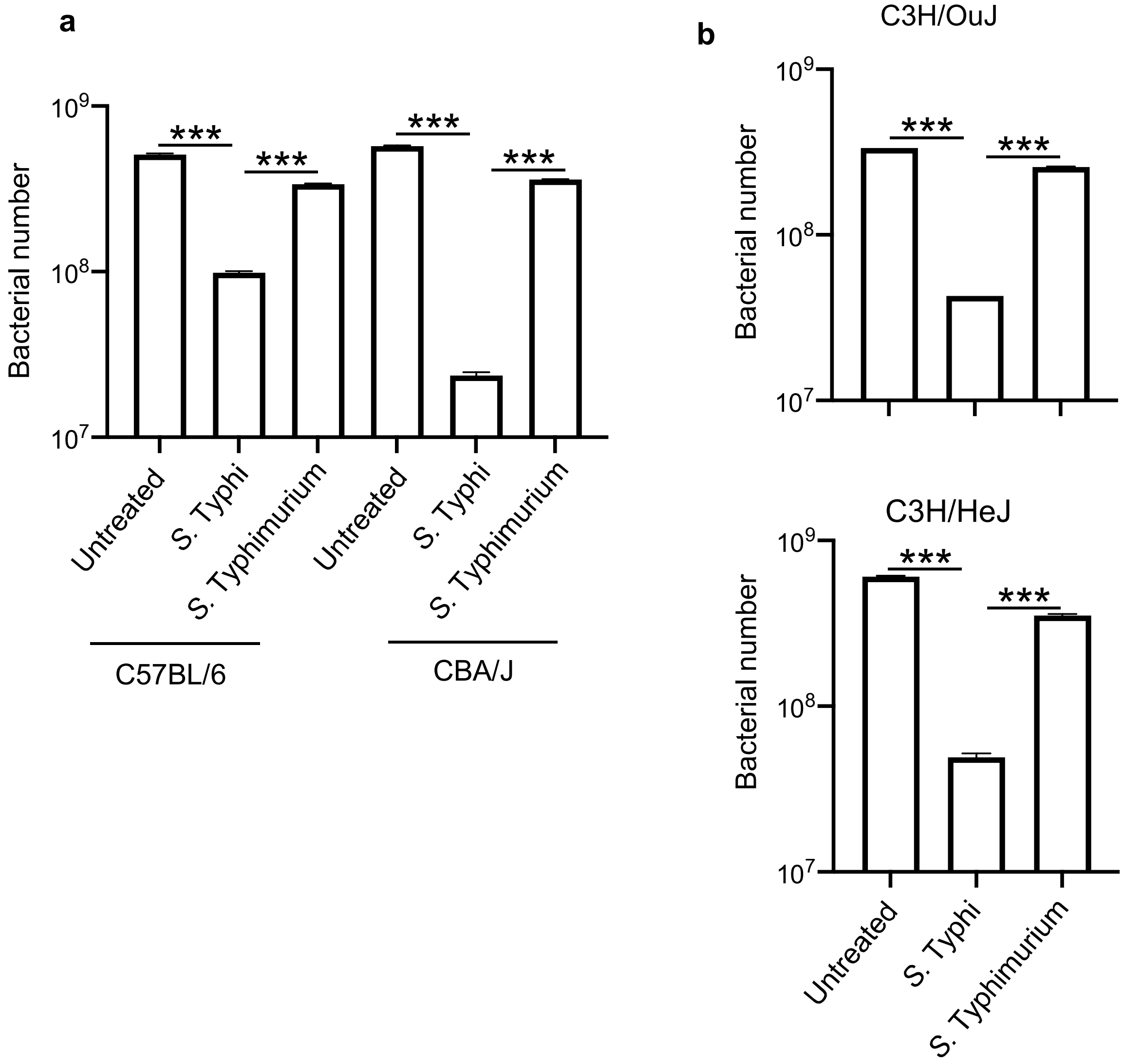


**Supplementary Figure 7: Cell-free supernatants from *S*. Typhi – infected macrophages inhibit growth of *Salmonella*.**

Peritoneal cells from different strains of mice were seeded in a 6 well plate ($4x10^6$ per well). 48 h later, cells were infected with *S.* Typhi or *S.* Typhimurium. Cell supernatants were collected at 2.5 h after infection and filtered through a 0.22 μ membrane. 10,000 *S.* Typhimurium were incubated with these supernatants at 37°C. Bacterial growth was determined after 9 h by measuring absorbance at 630 nm. Data representative of three independent experiments is shown as mean ± sem.. *** p<0.005.

#### Supplementary Figure 8

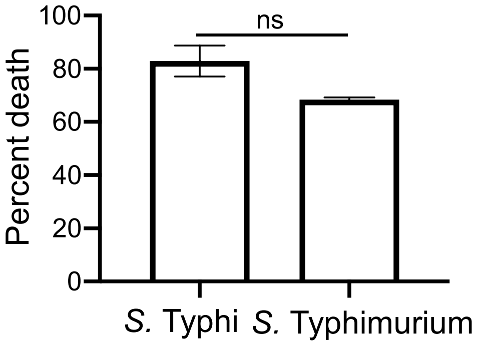


**Supplementary Fig. 8**: **Infection with *S*. Typhi or *S*. Typhimurium produces comaparable death in mouse macrophages**.

Resident peritoneal cells from C57BL/6 were seeded in a 6 well plate ($4x10^6$ per well). 48h later cells were infected with *S.* Typhi or *S.* Typhimurium for 2.5h. Cell death was determined by release of LDH in the supernatant. LDH released from cells treated with TritonX-100 (1%) was taken as 100% cell death. % lysis was calculated as (LDH released upon infection/LDH released by TritonX-100 treated cells) x 100. The results are representative of two independent experiments and are shown as mean ± sem. ns – not significant.

#### Supplementary Figure 9

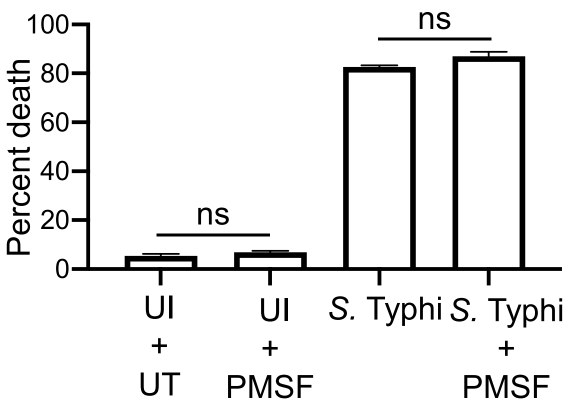


**Supplementary Fig. 9**: **PMSF does not affect *S*. Typhi – activated macrophage cell death**.

Resident peritoneal cells were pretreated with PMSF (100 µM) or with the vehicle (DMSO) and 30 min later, infected with *S.* Typhi for 2.5 h. Cell death was determined by release of LDH in the supernatant. The results are representative of two independent experiments and are shown as mean ± sem. UI **–** uninfected, UT–untreated, ns – not significant

**Supplementary Figure 10**

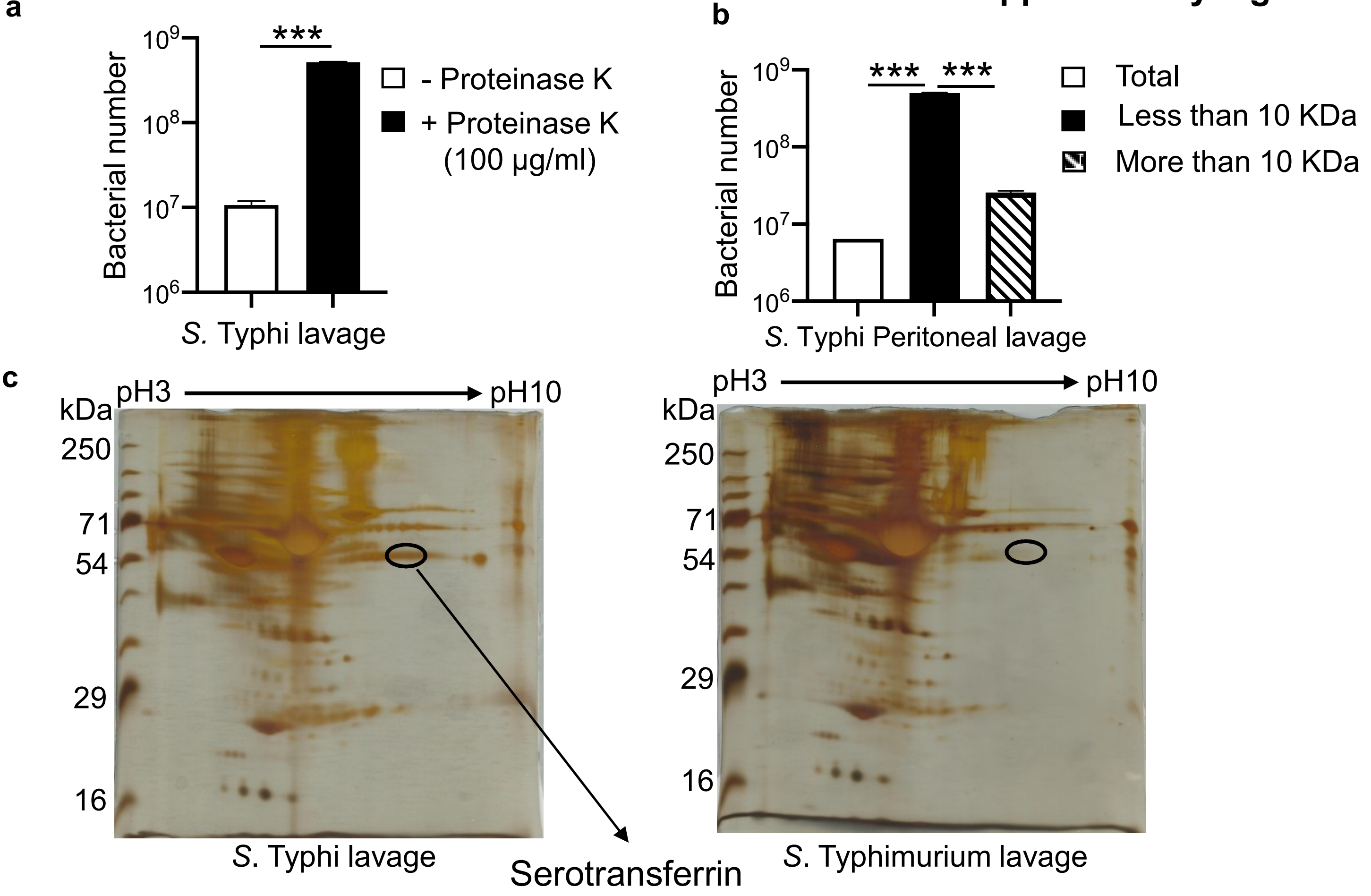


Table: Mass spectrometer-based protein identification

A

| | |
|---|---|
| **Protein accession number** | Q921I1 |
| **Protein name** | Serotransferrin |
| **Score** | 526.32 |
| **Coverage** | 27.55 |
| **Unique Peptides** | 17 |
| **PSMs** | 215 |

B

| **Peptide sequence** |
|---|
| IEcESAETTEDcIEK |
| NQQEGVcPEGSIDNSPVK |
| DGGGDVAFVK |
| cDEWSIISEGK |
| DQYELLcLDNTR |
| QEDFELLcPDGTR |
| GTDFQLNQLEGK |
| ASDTSITWNNLK |
| FDEFFSQGcAPGYEK |
| YLGAEYmQSVGNmR |
| LPEGTTPEK |
| GDVAFVK |
| DSAFGLLR |
| TSYPDcIK |
| EEYNGYTGAFR |
| TVLPPDGPR |
| ScHTGLGR |

**Supplementary Fig. 10**: **Cleaved serotransferrin is present in the peritoneal lavages of *S*. Typhi but not *S*. Typhimurium-infected mice.**

(**a**) Cell-free peritoneal lavage from *S.* Typhi - infected mice was treated with Proteinase K (100 µg/ml) for 1h. Antibacterial activity was determined by incubating 10,000 *S.* Typhimurium at 37ºC with undigested and Proteinase K - digested peritoneal lavage. Bacterial growth was determined by measuring absorbance at 630 nm. The results are representative of two independent experiments and are shown as mean ± sem. *** p value <0.005. (**b**) cell-free supernatants from uninfected and infected peritoneal cells were passed through a 10 kDa membrane and antibacterial activity was determined by incubating the retente and the flow-through with 10,000 CFU of *S.*Typhimurium. Bacterial growth was monitored at 630 nm. The results are from three independent experiments. (**c**) Peritoneal lavages collected after 4h of intraperitoneal infection with $5x10^6$ *S.* Typhimurium or *S.* Typhi were passed through 0.22 µ membrane and subjected to 2D gel electrophoresis. After silver-staining, a distinct spot present only in *S.* Typhi lavage was analyzed by mass spectrometry and identified as Serotransferrin. Table A depicts mascot score and coverage of the identified protein, and Table B shows unique peptides that matched with Serotransferrin.
